# Transformer with Convolution and Graph-Node co-embedding: An accurate and interpretable vision backbone for predicting gene expressions from local histopathological image

**DOI:** 10.1101/2023.05.28.542669

**Authors:** Xiao Xiao, Yan Kong, Zuoheng Wang, Hui Lu

**Affiliations:** State Key Laboratory of Microbial Metabolism, Joint International Research Laboratory of Metabolic and Developmental Sciences, Department of Bioinformatics and Biostatistics, School of Life Sciences and Biotechnology, Shanghai Jiao Tong University, Shanghai, China; SJTU-Yale Joint Center for Biostatistics and Data Science, National Center for Translational Medicine, Shanghai Jiao Tong University, Shanghai, China; Department of Biostatistics, Yale University, New Haven, CT, United States; Center for Biomedical Informatics, Shanghai Children’s Hospital, Shanghai, China

**Keywords:** deep learning, breast cancer, convolutional neural network, graph neural network, transformer, spatial transcriptomics

## Abstract

Inferring gene expressions from histopathological images has always been a fascinating but challenging task due to the huge differences between the two modal data. Previous works have used modified DenseNet121 to encode the local images and make gene expression predictions. And later works improved the prediction accuracy of gene expression by incorporating the coordinate information from images and using all spots in the tissue region as input. While these methods were limited in use due to model complexity, large demand on GPU memory, and insufficient encoding of local images, thus the results had low interpretability, relatively low accuracy, and over-smooth prediction of gene expression among neighbor spots. In this paper, we propose TCGN, (Transformer with Convolution and Graph-Node co-embedding method) for gene expression prediction from H&E stained pathological slide images. TCGN consists of convolutional layers, transformer encoders, and graph neural networks, and is the first to integrate these blocks in a general and interpretable computer vision backbone for histopathological image analysis. We trained TCGN and compared its performance with three existing methods on a publicly available spatial transcriptomic dataset. Even in the absence of the coordinates information and neighbor spots, TCGN still outperformed the existing methods by 5% and achieved 10 times higher prediction accuracy than the counterpart model. Besides its higher accuracy, our model is also small enough to be run on a personal computer and does not need complex building graph preprocessing compared to the existing methods. Moreover, TCGN is interpretable in recognizing special cell morphology and cell-cell interactions compared to models using all spots as input that are not interpretable. A more accurate omics information prediction from pathological images not only links genotypes to phenotypes so that we can predict more biomarkers that are expensive to test from histopathological images that are low-cost to obtain, but also provides a theoretical basis for future modeling of multi-modal data. Our results support that TCGN is a useful tool for inferring gene expressions from histopathological images and other potential histopathological image analysis studies.

**Highlights:**

1. First deep learning model to integrate CNN, GNN, and transformer for image analysis
2. An interpretable model that uses cell morphology and organizations to predict genes
3. Higher gene expression prediction accuracy without global information
4. Accurately predicted genes are related to immune escape and abnormal metabolism
5. Predict important biomarkers for breast cancer accurately from cheaper images

**Graphical abstract:** 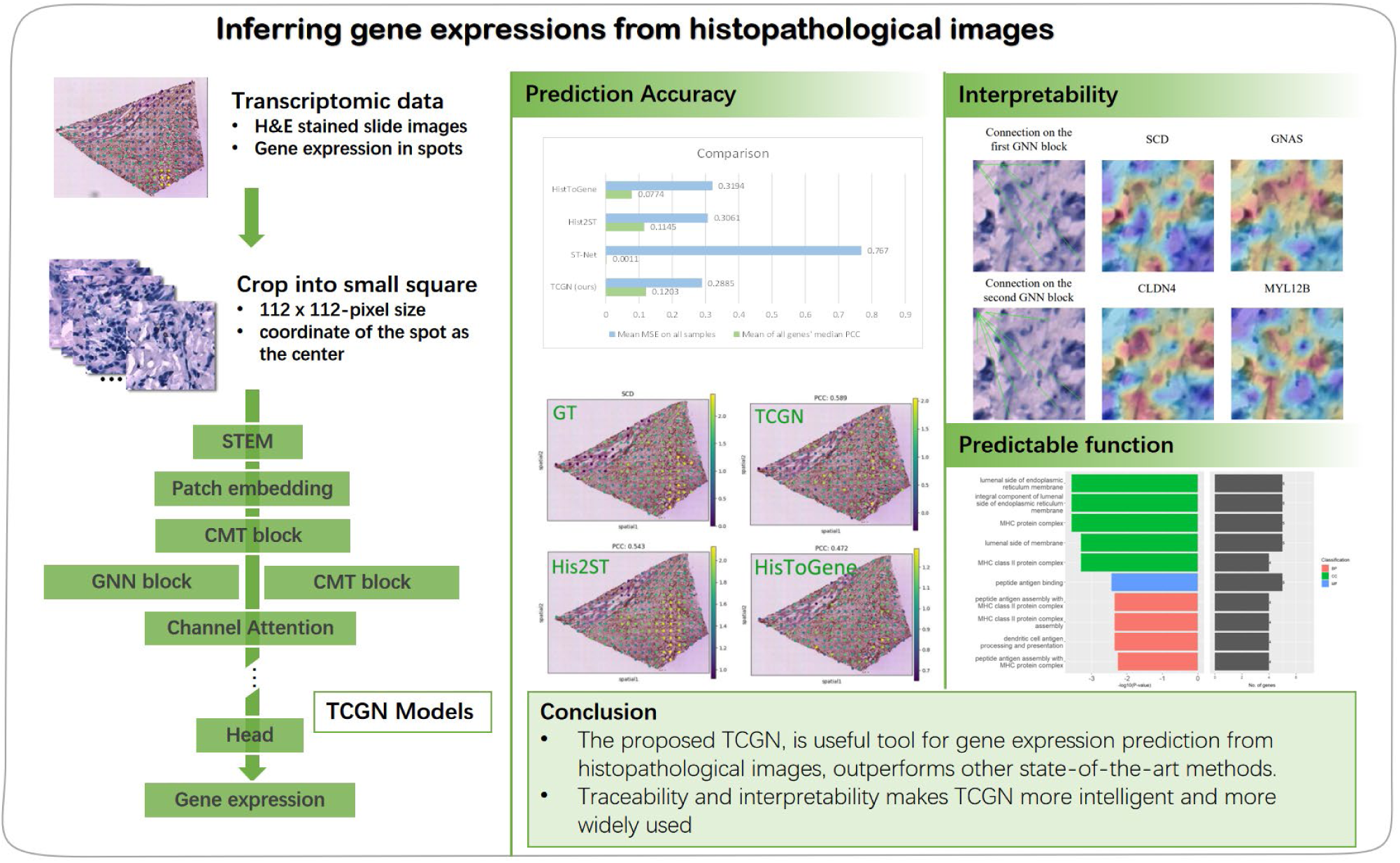

## 1. Introduction

Histopathological images, especially high-resolution microscopic images stained with hematoxylin-eosin, are important components for clinicians at the macro level to make cancer diagnoses and therapy decisions[1]. Digital pathological slide images together with molecular omics data have become essential support for oncology research[2–4]. At the molecular level, gene expression, mutation, and methylation levels have shown their ability to predict prognosis[5], identify biomarkers[6, 7], characterize cancer subtypes[8], and understand the mechanism of cancer[9, 10]. For biomedical images, some morphological features extracted from the pathological images are also believed to be related to the clinical outcomes of patients[11]. Emerging technologies and approaches have brought new opportunities to generate more accurate gene expression[12], mutation[13], and cell type annotations[14] at a single cell[15] or spatial resolution[16], which also brings the urgent need for linking molecular features and biomedical images.

Different modalities of multi-modal data are correlated with each other although they dissect tumors from different perspectives[17], and omics and pathological slides are no exceptions. The associations between phenotype and genotype have always been complicated but important problems for scientists. Of all these studies, predicting gene expressions from histopathological images is a very challenging task, if successful, it will broaden the applications of histopathology[18]. Since it is cheaper to obtain histopathological images than to sequence RNA expression and detect DNA mutations, if patients’ histopathological images can be used to predict the genotype of the patients with the help of explainable artificial intelligence, we will substantially reduce the cost of precision medicine and the workload of clinicians.

Previous studies have relied on statistical methods (e.g. linear regression), or traditional machine learning methods (e.g. random forest) to detect the relationship between hand-made H&E stained image features and gene expression features[3, 19]. However, their results were not as accurate as those based on recent deep-learning methods. With the wide application of deep learning methods in medical image analysis, such as image segmentaion[20, 21] and classification[4], many studies have demonstrated excellent performance of deep neural networks such as convolutional neural network (CNN) in building the connection between biomedical images and phenotypes such as microsatellite stability[4] and EGFR mutation status[22]. Despite the advantage, CNN has its limitation in dealing with complicated features such as graph-structured data. Meanwhile, thanks to the advance in mining graph data, graph-based deep learning model such as graph neural network (GNN) has become popular in histopathological images. Existing studies have shown that combining the utility of GNN and CNN can improve the performance of image analysis in prognostic prediction using multi-modal fusion techniques in gene expression prediction[23]. An essential step in the study of the relationships between gene expression and image features is to obtain cells with both single-cell gene expression and images to serve as ground truth. With the fast development of spatial transcriptomics[16], we can obtain image features for several cells and the corresponding gene expression profiles from a relatively small area, which allows us to conduct the association study of genotypes and phenotypes through deep learning[3].

There are currently three studies that performed gene expression prediction from histopathological images in spatial transcriptomics. ST-Net[3] used only the local spot image in spatial transcriptomics, while HisToGene[24] and Hist2ST[25] treated all spots in the tissue section and their coordinate information as input to predict gene expressions of all spots. The prediction accuracy of ST-Net is low, with the median Pearson Correlation Coefficient (PCC) of all the predicted highly variable genes of 0.0011, so the number of statistically significant predictable genes is very small. Due to the incorporation of information from all the spot images in one tissue section, Hist2ST and HisToGene outperformed ST-Net to a large extent and achieved the median PCC of all the highly variable genes of 0.1245 and 0.0774, respectively. Although Hist2ST and HisToGene achieved higher prediction accuracy by utilizing coordinate information and self-attention, there is still room for improvement in their methods and results. First, Hist2ST encoded the image data through convolutional layers, while they were not interconnected with the transformer encoder blocks. So it would only learn image features extracted from convolutional layers and then assign attention to the extracted features, but more complex features that the convolutional layers do not extract were not able to be used. The gene expression prediction accuracy can be improved by fully utilizing local image features. Second, by adopting the spatial graph convolutional neural network to aggregate features between neighbor spots, Hist2ST tended to over smooth the predictions at neighborhood spots. However, the same gene can have very different expressions at neighboring spots. Third, the variation of gene expression prediction can be large among different sections in the leave-one-out cross-validation step, as the prediction accuracy of the same gene varies largely among different sections. Finally, using all the spots as input in deep learning models to predict gene expression for each spot takes too much GPU memory to run on a personal computer and can suffer from interpretability, because it is difficult to quantify the contribution of each pixel to the gene expression prediction. Thus, it is important to develop an interpretable deep learning model that can fully utilize local histopathological image data without global information to predict the molecular signature and perform other histopathological image analysis tasks.

In this article, we developed TCGN-<u>T</u>ransformer with <u>C</u>onvolution and <u>G</u>raph-<u>N</u>ode co-embedding to predict gene expression from histopathological images in spatial transcriptomics using only the local spot image. The proposed TCGN is a novel deep-learning model which takes the advantage of CNN, the transformer encoder, and GNN in a single computer vision backbone for biomedical image processing. Our model can extract local features using CNN, learn representations using the transformer, and analyze graph-structure features using GNN. We demonstrated that TCGN has an improved performance in comparison with several commonly used methods in predicting gene expression from histopathological images with smaller model size and without coordinates information and neighbor spots. Functional analysis results of high-accuracy genes predicted from histopathological images also showed the power of our model in predicting important breast cancer biomarkers and proved the interpretability of our proposed method.

## 2. Material and methods

### 2.1 Datasets and pre-processing

In this study, we used the public human HER2-positive breast tumor dataset (HER2+)[26] to evaluate the performance of TCGN. This dataset contains the spot image, spot coordinate, and the corresponding gene expression at each spot. The data was measured by the spatial transcriptomics technique and contains 36 tissue sections from 8 patients who were diagnosed with breast cancer and tested positive for the human epidermal growth factor receptor 2 (HER2) protein. In our study, the spot images were treated as the input, and the corresponding gene expression in this spot as the label.

The preprocessing workflow was also similar to the previous works[24, 25]. For each spot in the dataset, we cut out a small square piece with a 112×112-pixel size with the coordinate of the spot as the center. Then we normalized the gene expression levels in each spot with library size normalization and log transformation using the python package scanpy[27]. Specifically, the count of each gene in the spot was divided by the sum of all the counts in this spot, then multiplied by a scale factor of 1,000,000 and log-normalized as ln(1 + *x*), where x was the gene count divided by the total count and multiplied by 1,000,000. We further excluded sections that had less than 180 spots in this section (A1, H1, H2, H3) after normalization. This resulted in 32 sections. Then, we selected the top 1000 genes with the largest standard deviation and excluded genes that expressed in less than 1000 spots. After filtering, there were 785 genes that remained for analysis.

### 2.2 Model Architecture

The whole architecture of the proposed model is shown in **Figure 1**. The main component includes the convolutional layer, the transformer[28] with position embedding bias[29] section, and the GNN section. The input image with a size of 224×224 pixels first goes through a stack of convolutional layers arranged like ConvNext[30] for the model to extract its local and primary features. Then it is reshaped by the patch embedding layer and processed by the CNN meet vision transformer (CMT) blocks. While CMT uses the self-attention mechanism to learn complicated features, the GNN block also uses the features processed by the previous CMT blocks to learn the graph-structured cell organization features in the image. After extracting different kinds of features and using channel attention to integrate these features, we used a 1 × 1 convolutional layer, batch normalization layer, GELU activation function, and average pooling to aggregate all features and make the final prediction.

**Figure 1.**
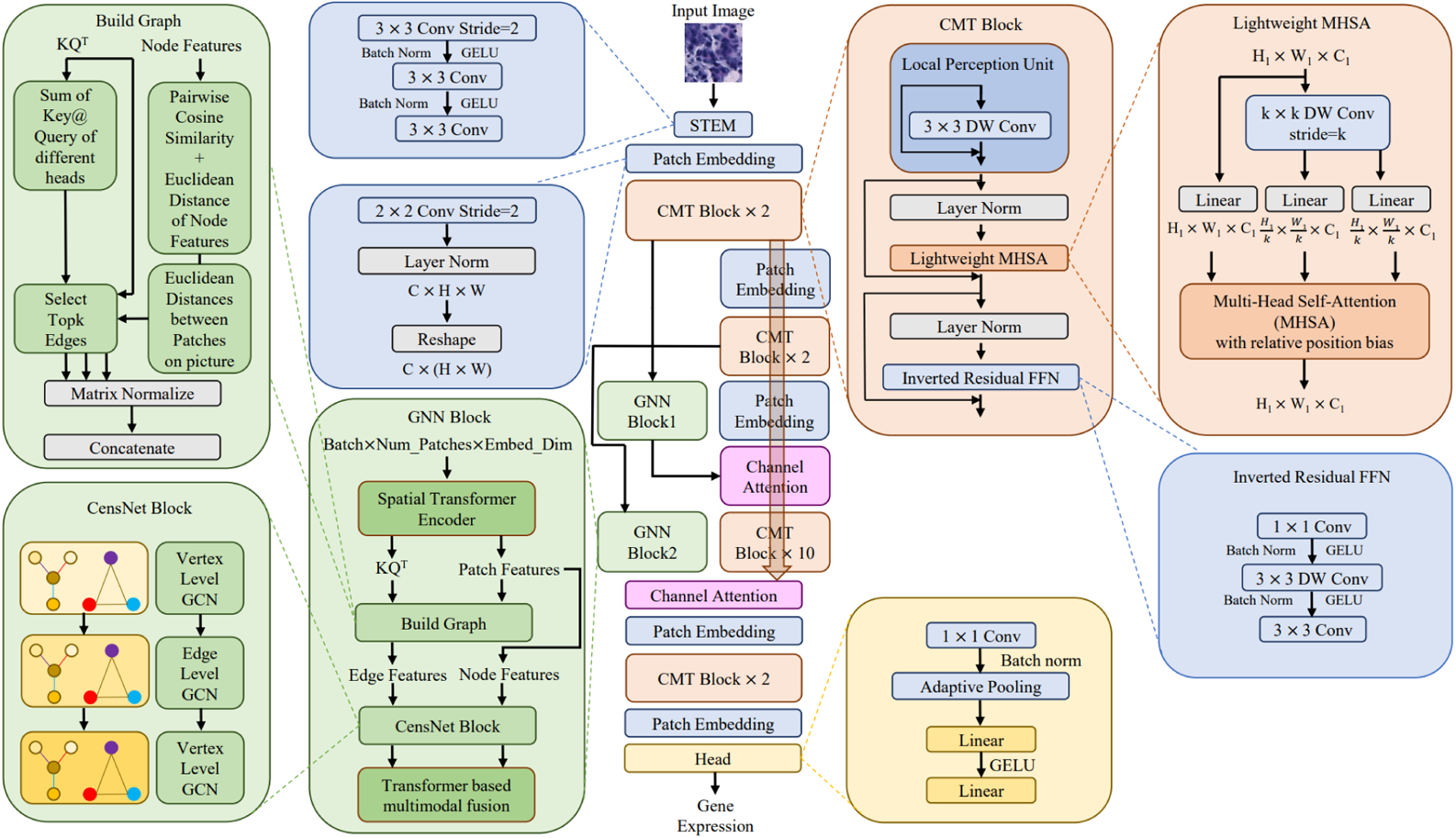
The architecture of TCGN. It consists of the convolutional layer, transformer encoder, and graph neural network block. The features will first go through some convolutional layers and transformer encoders to learn primary representations and local and simple features. Then it will go through two GNN blocks that are not interrelated. Then it will further go through other CMT blocks and finally go through a prediction head to make gene expression prediction.

#### 2.2.1 CMT block

We introduce the CMT block[29] instead of the vision transformer[31] encoder in our model to prevent potential problems such as large dataset requirements, 2D information loss in 1D linear transformation within self-attention, insufficient encoding of local features, and damaging biomedical images’ translation invariance[29]. With well-arranged architecture, the CMT block can successfully combine the convolutional layer with the transformer and ingeniously optimize the self-attention mechanism using a small size of parameters. The CMT block consists of three parts, including the local perception unit (LPU), the lightweight multi-head self-attention (LMHSA), and the inverted residual feed-forward network (IRFFN), and they are connected through layer norm and residual block. The details of these architectures and how they are designed will be illustrated below. Taking *X* as the input, the overall procedure of the CMT block can be explained by the following formula:

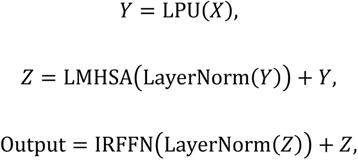

Where *Y* is the output of the LPU block and *Z* is the output of the LMHSA block.

##### 2.2.1.1 Local perception unit (LPU) and inverted residual FFN (IRFFN)

LPU is designed to capture local features and retain translation invariances in the CMT block, which could overcome the traditional vision transformer’s limitation of insufficient encoding of local features as well as loss of translational invariance caused by asymmetric position encoding[29].

In order to prevent local information loss, LPU is used to capture local features. LPU consists of a depth-wise convolution block and a residual block. By treating each feature independently through depth-wise convolution, we utilize global information without damaging the translation invariance. The depth-wise convolution manipulation is also much faster than the traditional convolutional layer, thus speeding up the computation. With the input tensor *X* of a dimension of *H* × *W* × *d*, where *H* and *W* are the height and width of the image and *d* is the dimension of features, LPU is defined as:

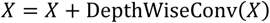

The IRFFN block’s function is the same as that of the feed-forward network (FFN) block in the transformer encoder. Similar to LPU, the IRFFN block uses the convolutional layers to replace the multi-layer perceptron (two linear layers) in order to retain translation invariance and 2D structure. Following the guidance that FFN has the same number of nodes at the input and output side and another number of nodes in the middle layer, IRFFN is defined by a 3 × 3 convolutional layer covered by two 1 × 1 convolutional layers. Notably, the GELU activation function and the residual block are used the same as the transformer encoder to avoid vanishing and exploding gradients.

##### 2.2.1.2 Lightweight MHSA

The lightweight MHSA is designed to solve the problem of losing 2D information in the Vision Transformer, where the key (K), query (Q), and value (V) in the self-attention block are simply generated by linear transformations. However, these linear transformations are manipulations on 1D data which may not be appropriate for patches’ 2D structure features and images’ translation invariance.

Unlike ViT, the CMT block does not have operations that directly transform a reshaped tensor with a dimension of *C*_1_ × *n* to a tensor with a dimension of *C*_2_ × *n*, where *n* = *W* × *H*. Instead, the CMT block produces K and V through *k* × *k* depth-wise convolutional layers plus a linear transformation so that the 2D information would not be lost. To learn the position information negatively influenced by the tensor reshaping operation, the CMT block also adds a learnable variable-relative position bias (B) to the self-attention block. The whole procedure of the lightweight MHSA is defined below:

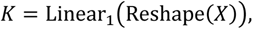

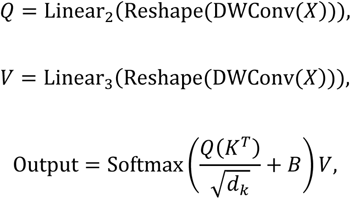

where 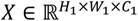 is the input, DWConv refers to *k* × *k* depth-wise convolution with stride k, B donates the trainable relative position bias parameter, and *d_k_* is the embedding dimension of K.

#### 2.2.2 GNN block

The GNN block is designed to extract the graph-structure features of the input histopathological image. It is also responsible for predicting how these graph-structure features will interact with the features that the CNN and the transformer encoder extract from the image. Generally, the architecture of the GNN block consists of the connection analyzer, the edge features encoder, the CensNet block, and a multimodal features fusion encoder (**Figure 2A**).

**Figure 2.**
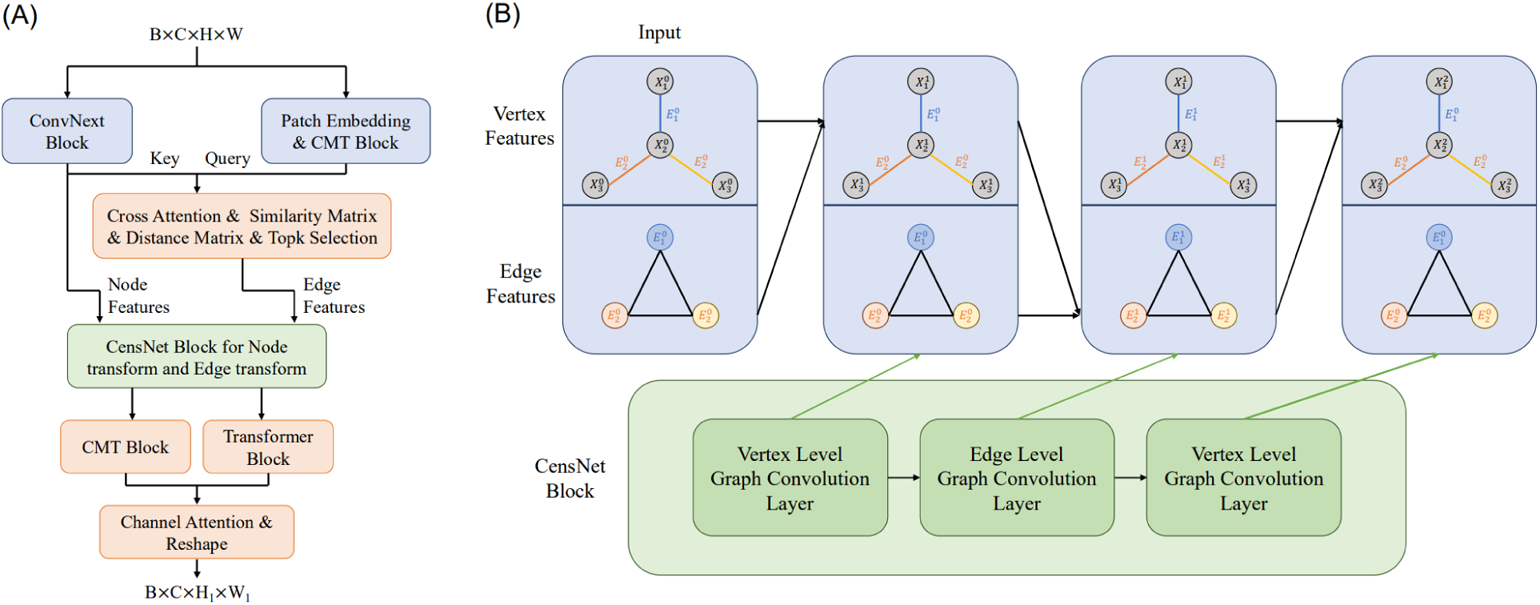
The architecture of the GNN block. **A.** The overall procedure of the GNN block, which consists of the connection analyzer, the CensNet block, and the fusion block. **B.** The architecture of the CensNet block, which used the graph convolution with node-edge switch to deal with multi-dimensional edge features and multi-dimensional node features.

##### 2.2.2.1 Connection analyzer

We design our connection analyzer in order to analyze the complex interactions between patches in the input image, which contains interactions between cell nuclei and cytoplasm. In the GNN block, patches will first go through a connection analyzer to analyze the complex interactions between patches and cells in the input image. The connections between two patches depend on the features and the distance of each other. A ConvNext block is used to extract the texture features of the image which is used in training steps to automatically learn how patches with certain morphological features would interact with each other. A CMT block is applied to introduce the distance/position information to the connection analyzer and to extract some texture features within the patch. Finally, we use the cross-attention block after the ConvNext block and the CMT block to integrate the features information and the position information for building the graph. To be more specific, the input tensor with shape batch_size × channel × height × width will first go through a ConvNext block[30] for the model to extract its local texture features. At the same time, the input will also go through a CMT block for the model to learn the representations. Then a multi-head cross-attention block is used where the features generated by the ConvNext block are treated as the key and the features generated by CMT are treated as the query. The whole procedure of the connection analyzer is defined below:

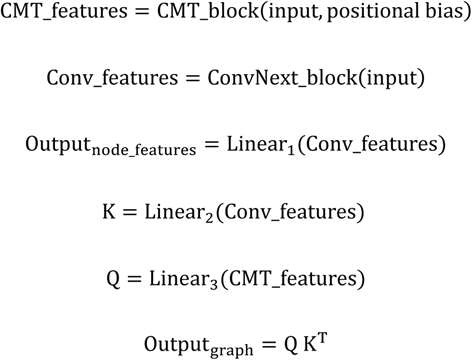

##### 2.2.2.2 Edge features encoder and the top K selection

In order to let our model better analyze the connections between patches, we additionally add the similarity measurements to the edge features generated by the connection analyzer. The intuition is that the contribution of cells with the same features to the overall gene expression may be the same and can be aggregated by the graph convolutional neural network. So similar to Vision GNN[32], we add the L2 distance and cosine distance and perform a transformation to these two metrics so that the values would be ranged from 0 to 1 and larger values mean closer connections, which is the case in graph convolution operations.

The detailed formula of the edge features between node/patch *i* and *j* is as below:

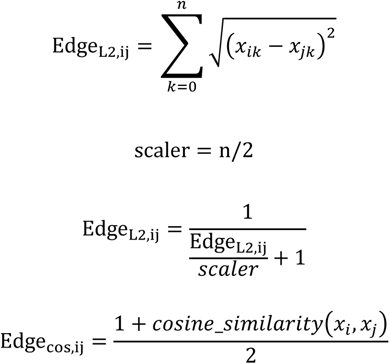

Where n is the number of dimensions for each node/patch. In addition, the calculation of L2 distance and Cosine distance requires the gradient.

Besides the Cosine distance and L2 distance, we also add the real distance between patches in order to tell GNN to take more attention to the position information. And the normalization method for this value is as below:

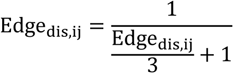

Contrary to the cosine distance calculation, as the distances between each patch are the same for different input images, this calculation does not require gradients.

After building the graph according to cross-attention and our hand-make features, these edge features are concatenated as a complete description of edge features. Due to the inequality of contributions of patches and connections for gene expression levels, and for reduction of computation, we sort and select top K connections based on the node adjacent matrix. To be more specific, we only select the top number_of_patches × topk edges in the adjacent matrix according to the values of the sum of all the corresponding edge features. We set the edge features that are not selected to the value of 0. As shown in most previous works[23, 33] using spatial cell graphs, the K was set to 4.

##### 2.2.2.3 Preliminaries for the CensNet Block

We assume that we have a directed graph G input with multi-dimensional edge E features and multi-dimensional node V generated by our connection analyzer. We define the input elements of our CensNet block as below.

- Let 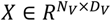 be the node feature matrix, where *N_v_* refers to the number of nodes in the graph, which is (224/16) × (224/16) = 196 in our model, and *D_v_* refers to the number of dimensions of node features
- Let 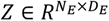 be the edge feature matrix, where N_E_ donates the number of edges, which is topk × number_of_nodes = 4 × 196 = 784 in our model, and D_E_ refers to the number of dimensions of the edge, which is number_of_head_in_cross_attention + 3 in our model.
- Let 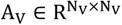 be the node adjacent matrix. The diagonal elements of the matrix may be set to zero according to the output of cross-attention, and the values in this matrix are continuous instead of only 0 or 1.
- Let 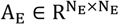 be the edge adjacent matrix. The diagonal elements of the matrix are always non-zero, and the values in this matrix are discrete.
- Let 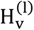 be the l-th hidden layer of the vertex features, and let 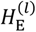 be the *l*-th hidden layer of the edge features. Specifically, 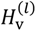 equals to *X* and 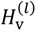 equals to *Z* at the 0-th layer.

##### 2.2.2.4 Building edge adjacent matrix and transition matrix from edge features

Unlike ordinary graph convolutional neural networks, the CensNet we use in our model utilizes multi-dimensional edge features and multi-dimensional node features and adopts the node-edge switch to learn the graph features from different perspectives. To achieve that, the edge adjacent matrix and the transition matrix between the node adjacent matrix and the edge adjacent matrix are in need.

The edge adjacent matrix 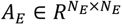. There is a connection between *edge_i_* and *edge_j_* if start_i_ == start_j_ or start_i_ == end_j_ or end_i_ == start_j_ or end_i_ == end_j_, where start_i_ and end_i_ refer to the starting vertex and ending vertex of edge_i_, respectively. If there is a connection between edge_i_ and edge_j_, then A_E,ij_ = 1, otherwise *A_E_*_,*ij*_ = 0. And Finally, the edge adjacent matrix was Laplacian normalized with the formula A_E_ = D^−0.5^A_E_D^−0.5^, where D is the degree matrix of A_E_. As a result, the diagonal elements of A_E_ are always non-zero.

The transition matrix 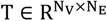. T_i,m_ = 1 if node_i_ is the starting point or ending point of edge_m_ and all other elements in the matrix are 0.

##### 2.2.2.5 CensNet block

In this work, we choose to use CensNet[34] as the graph neural network in our model to extract the graph features and aggregate information between different patches. Unlike the traditional graph convolutional neural network, the CensNet innovatively uses the node-edge switch to make use of the multi-dimensional features in the heterogeneous graph. Moreover, the CensNet block can analyze the interactions between the node features. And the edge features will be analyzed by using both the node features and the edge features to make a transformation for only the node features or the edge features. Notably, if we simply use a traditional graph convolutional neural network, the GNN block will just become another transformer block, because a simple graph neural network is just another form of attention mechanism[35].

The CensNet block consists of the node transformation layer and the edge transformation layer. As most graph convolutional neural networks cannot go deep[36], otherwise the output would be too smooth, the CensNet block in our work only consists of two node transformation layers and one edge transformation layer. A simple illustration of the CensNet block is shown below in **Figure 2B**.

For the node transformation layer, the forward propagation rule from *l*-th node hidden layer to *l* + 1-th node hidden layer is:

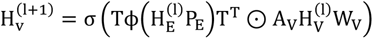

This formula can be viewed as using edge features to “adjust” the node adjacent matrix and performing a graph convolution based on the “adjusted” node adjacent matrix. In this formula, 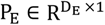 is a learnable parameter, and is used to make a linear transformation for the edge features.

And ϕ(Input) donates the diagonal embedding operation, which means creating a matrix where the diagonal elements are the input 1-dimensional vector and all other elements are 0. After transforming a multi-dimensional edge feature to a single value with the help of P_E_, we then use the transition matrix T to add this information to the node adjacent matrix to form an “adjusted” node adjacent matrix through Hadamard product ⊙. Then, we use W_V_, a learnable parameter to transform the node features, and used ϕ, an activation function, to add no-linearity and complete the graph convolution process.

Similarly, for the edge transformation layer, the forward propagation rule from *l*-th edge hidden layer to *l* + 1-th edge hidden layer is:

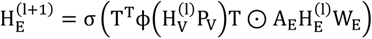

The method is similar to the node transformation layer: use P_V_ to learn how node features would influence the edge adjacent matrix, and perform a graph convolution based on the “adjusted” edge adjacent matrix.

Because the interaction between patches/cells in the picture is very complex and can be affected by multiple factors, the idea of using node features to modify the edge connection and using edge features to modify the node connection is very helpful for our model to further explore what the connection is and how to utilize the connection. This is also the reason why we added the distance and the similarity metric between patches to edge features, as features and distance together make a more complete connection strength.

##### 2.2.2.6 Multimodal fusion of graph-structure features and patch features

We use the transformer encoder, alignment between node and edge features through the transition matrix, and channel attention to further encode and fuse the node features and the edge features. The node and edge features will first be encoded by the transformer-based CMT block and original transformer block for the model to capture the long-distance dependencies that may not be able to be captured by simple graph convolutional neural networks that are not deep enough, and replace the graph pooling that may lose certain information and cause the over-smooth problem. This further encoding process is also beneficial for the later multi-modal fusion[37]. Then after primarily encoded by the transformer block, we multiply the edge features matrix 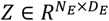 by the transition matrix 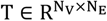 so that the edge feature matrix 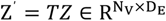 and the node feature matrix 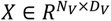 can be aligned and shared the same first-dimension number *N_v_*. At last, we use channel attention[38] to integrate the edge features and the node features of the corresponding node. The detail of this process can be described below:

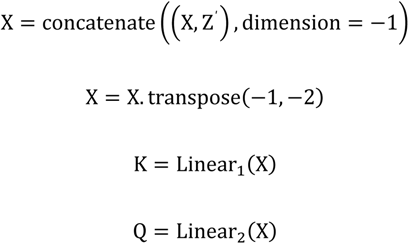

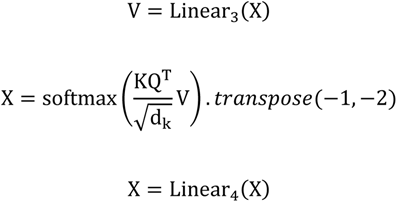

Where Linear_1_, Linear_2_, Linear_3_ are linear transforms from 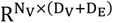 to 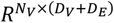, Linear_4_is the linear transform from 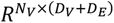 to 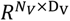, d_k_is the number of the last dimension of the transposed input, and transpose means to swap between two dimensions within a tensor.

#### 2.2.3 Synthesize different kinds of features

For the combination of the GNN block and the CMT block, we concatenate the features of the same patch in the last dimension and performed channel attention for each patch, so that the model can learn how these features will interact and pay attention to the features that make more contributions to the final gene expression prediction.

Another problem is that we want the model to extract the graph-structure features at different stages of the model so that we will have the node features in the graph with local features and with higher-level CNN features, but the GNN cannot go deep otherwise the output will be too smooth. So instead of letting the features pass through all the blocks in our model one by one, we integrate the output of the first GNN block and the CMT block after the second GNN block receives its input from the previous CMT block (**Figure 1**). In this way, the second GNN block will not receive its input processed by the first GNN block, and there would not be an over-smooth problem while extracting graph structure at different levels.

### 2.3 Experiment

We test our model and previous models (Hist2ST, HisToGene, ST-Net) on the HER2-positive breast tumor dataset and compared their performance. The input of our model is simply the RGB image, which is firstly resized from size 112 × 112 to size 224 × 224. Then we scaled all the values by dividing them by 255 and normalized the values of the RGB picture to the mean of (0.485, 0.456, 0.406) with the standard deviation of (0.229, 0.224, 0.225), which is the default parameter of the deep learning package timm[39]. In order to enhance the model’s generalization ability and prevent overfitting, we augmented our dataset when training by randomly rotating the image with ±180^°^, randomly make a horizontal flip and vertical flip with a probability of 0.5. For the output of our model, which is a tensor with shape batch_size × 785, we used the mean square error between the prediction and the ground truth as the loss function.

#### 2.3.1 Hyperparameter setting

The TCGN is implemented in Python 3.8 and Pytorch 1.13.1. For the CMT block, we set the CMT block’s hyperparameter of the number of heads, depth, and width to the default hyperparameter of CMT-Ti[29]. For the GNN block, the top_k is set to 4 for the connection analyzer and the dimension of the hidden layer is set to 92 for both the first and the second GNN block. For the prediction head, the number of channels for the last 1 × 1 convolutional layer is set to 1280, and the number of dimensions for the last two linear layers is 5120 and 785. The optimizer we used in our study was the adam[40], and the learning rate is set to 0.00001 with the coefficients used for computing running averages of gradient and its square set to 0.9 and 0.999. The model we save is the one that had the highest median PCC on the validation set among the models in different epochs. For more details, please refer to the released code on GitHub. Besides, all results reported in this article were conducted on CentOS Linux release 7.7.1908 with Intel Xeon Scalable Cascade Lake 8168(2.7GHz, 24 cores) and NVIDIA Tesla V100. For comparison, we run the code of previous works with the same hardware as above and the default parameters in the released version of their codes on GitHub. In addition, the comparison of model input type, number of parameters, model size, and training hyperparameters between 4 works are listed below in **Table 1**.

**Table 1.**
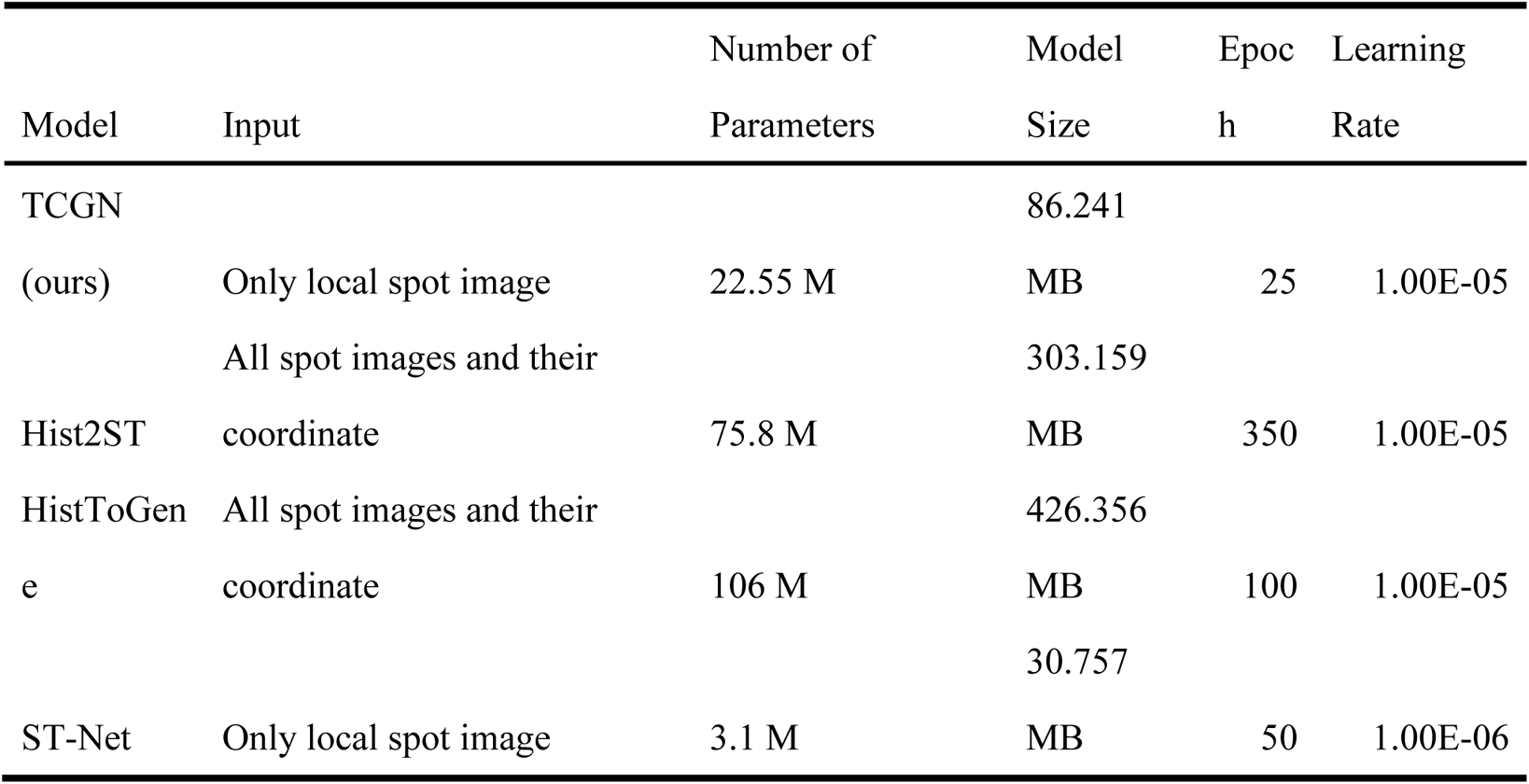
The comparison of the input size, the number of parameters, the model size, and the training hyperparameters between our work and the previous 3 works.

#### 2.3.2 Evaluation metrics

##### 2.3.2.1 Pearson correlation coefficients

For each section, we have a vector of X_predict_ and X_truth_ with the length of the number of spots in this section for every gene. So we evaluate our model and previous works through Pearson Correlation Coefficient (PCC), which is the same as the last 3 related works. To demonstrate the overall performance, the mean and median PCC across all sections in the 32 cross-validations for every 785 genes is also calculated.

The formula of the PCC is shown below:

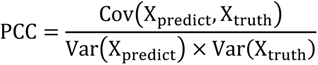

Where Cov refers to the covariance and Var donates the variance.

##### 2.3.2.2 Mean square error

Besides PCC, the mean square error (MSE) is also a very common metric for regression problems in deep learning, which directly measures the difference between the prediction and the ground truth. And the MSE loss is also the loss function in our work and previous works (ST-Net, HisToGene, and Hist2ST), which was used as the metric to optimize the deep learning models. As a result, we also evaluated our model and compared our model with previous models using the MSE loss metric.

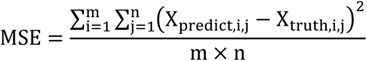

Where m is the number of spots in this section and n is the number of genes, which is 785. X_predict,i,j_refers to the predicted gene expression level of gene j in the spot i and so does X_truth,i,j_.

## 3. Result

### 3.1 Prediction Accuracy

We evaluated our model on the HER2-positive breast tumor dataset through leave-one-out cross-validation on 32 selected samples and compared it with 3 existing methods. TCGN achieved the best prediction accuracy based on both the PCC and MSE metrics, and more stable prediction among different samples in cross-validation without over-smoothness.

The detailed comparison of the PCC of all 785 genes on 32 different samples was demonstrated in **Figure 3A**. Our model achieved the best accuracy in terms of the mean of the median PCC of all genes, 5% higher than His2ST, the second-best model (Table 2). Compared to ST-Net which had the same input type as our model, TCGN achieved about 10 times higher median PCC. In addition, as shown in Figure 3A, although the interquartile ranges of the “mean” and the “median” boxplots are almost the same, our model had more genes whose values are higher than the third quartile plus 1.5 times of the interquartile range, suggesting that TCGN is able to predict more genes with high accuracy. Although the median PCC of TCGN on some samples was slightly lower than that of Hist2ST (Figure 3A), our model still achieved a better prediction accuracy in terms of median PCC and mean PCC of all genes across samples (Table 2).

**Figure 3.**
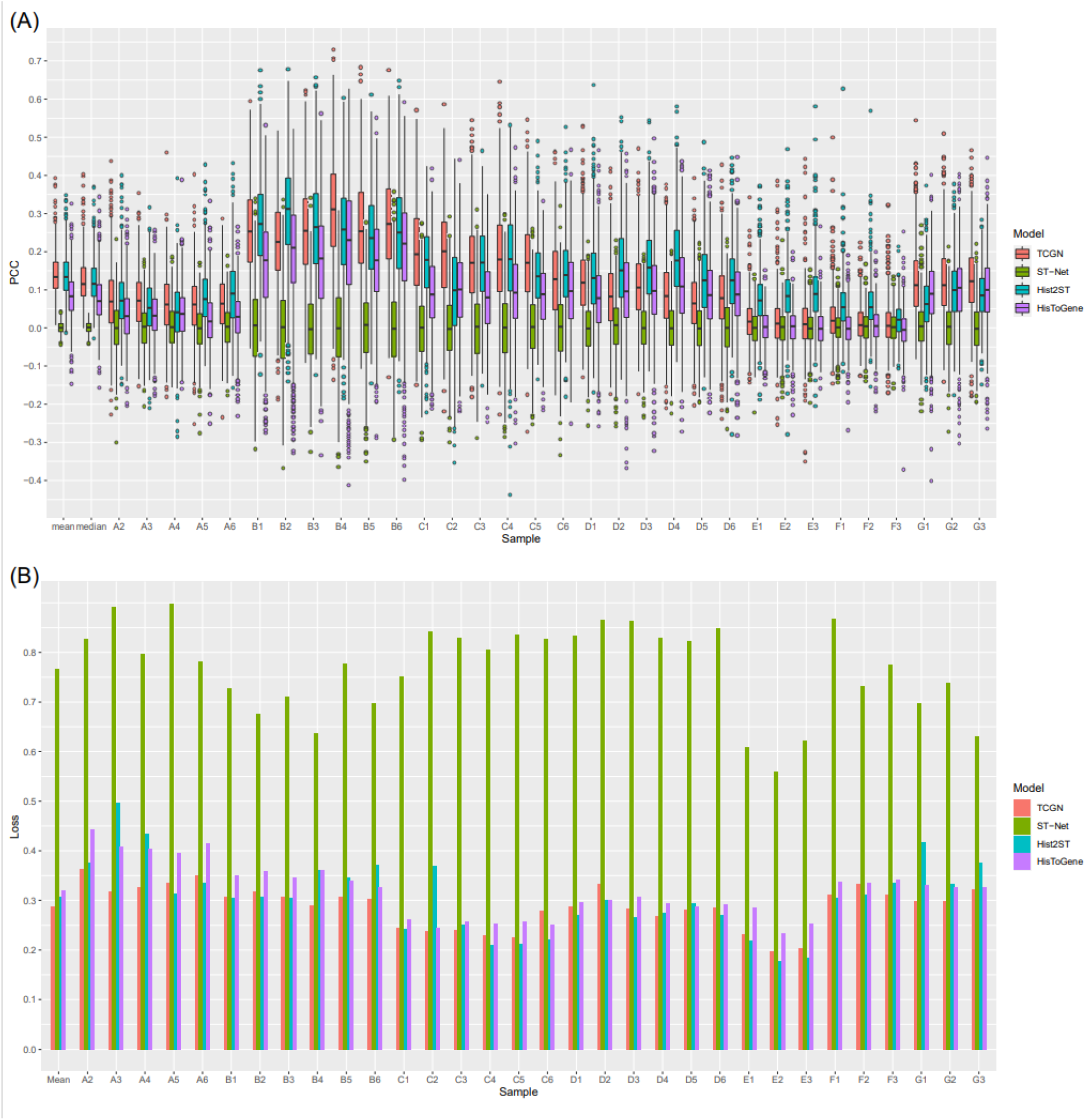
Comparison of our model with 3 previous models. **A.** The comparison of our model and the previous 3 models which were evaluated by the PCC of predicted gene expression and observed gene expression on 32 leave-one-out cross-validations and their mean and median. **B.** The comparison of our model and the previous 3 models in terms of the MSE loss on the validation set of the 32 leave-one-out cross-validations and their mean values.

**Table 2.** The comparison of the statistics of the MSE loss and the PCC metrics, as well as the standard errors of the MSE loss among different samples of the 4 models.

| Model | Mean of all genes' median<br>PCC | Mean MSE on all<br>samples | Standard error of all<br>MSE |
| --- | --- | --- | --- |
| TCGN<br>(ours) | 0.1303 | 0.2885 | 0.0412 |
| ST-Net | 0.0011 | 0.767 | 0.0877 |
| Hist2ST | 0.1245 | 0.3061 | 0.071 |
| HistToGene | 0.0774 | 0.3194 | 0.0531 |

To evaluate the overall performance of gene expression from the histopathological images, we evaluated and compared the MSE loss of the 4 models (Figure 3B). Our model had the lowest mean MSE loss across all samples, 6% lower than His2ST, the second-best model (Table 2), and only half of the MSE loss of ST-Net which used the same input as ours.

In order to assess the stability among different runs in cross-validation, we compared the variation of the MSE loss across samples, The results showed that TCGN had the smallest standard deviation of the MSE loss (Table 2), suggesting that our model can predict the genes in a more stable manner.

### 3.2 Ablation study

We performed an ablation study to understand the contribution of each block to the overall model. We used the median PCC for all 785 genes on sample A2 (fold 0) as the metric in the ablation study, and the optimizer and hyperparameters were the same as those in the assessment of TCGN. The ablation results were shown in **Figure 4**. We can see that adding the transformer encoder to the convolutional neural network (the CMT block) made the largest contribution to the model such that TCGN outperformed ST-Net (CNN) and the vision transformer. This has been observed in previous studies, such as Hist2ST, where the removal of the transformer module would decrease the prediction accuracy by 11% and has the largest influence on other modules in Hist2ST. Besides the CMT block, the GNN block also brought in a significant improvement of 4% to the model, suggesting that the graph-structured features from the histopathological image help characterize the tissue. In addition, a similar amount of improvement was also reported in a previous study[23], where the improvement was about 5%. For the channel attention block, if we replaced it with one linear layer, the median PCC on sample A2 decreased by 1%, showing the advantage of channel attention in fusing different kinds of feature information.

**Figure 4.**
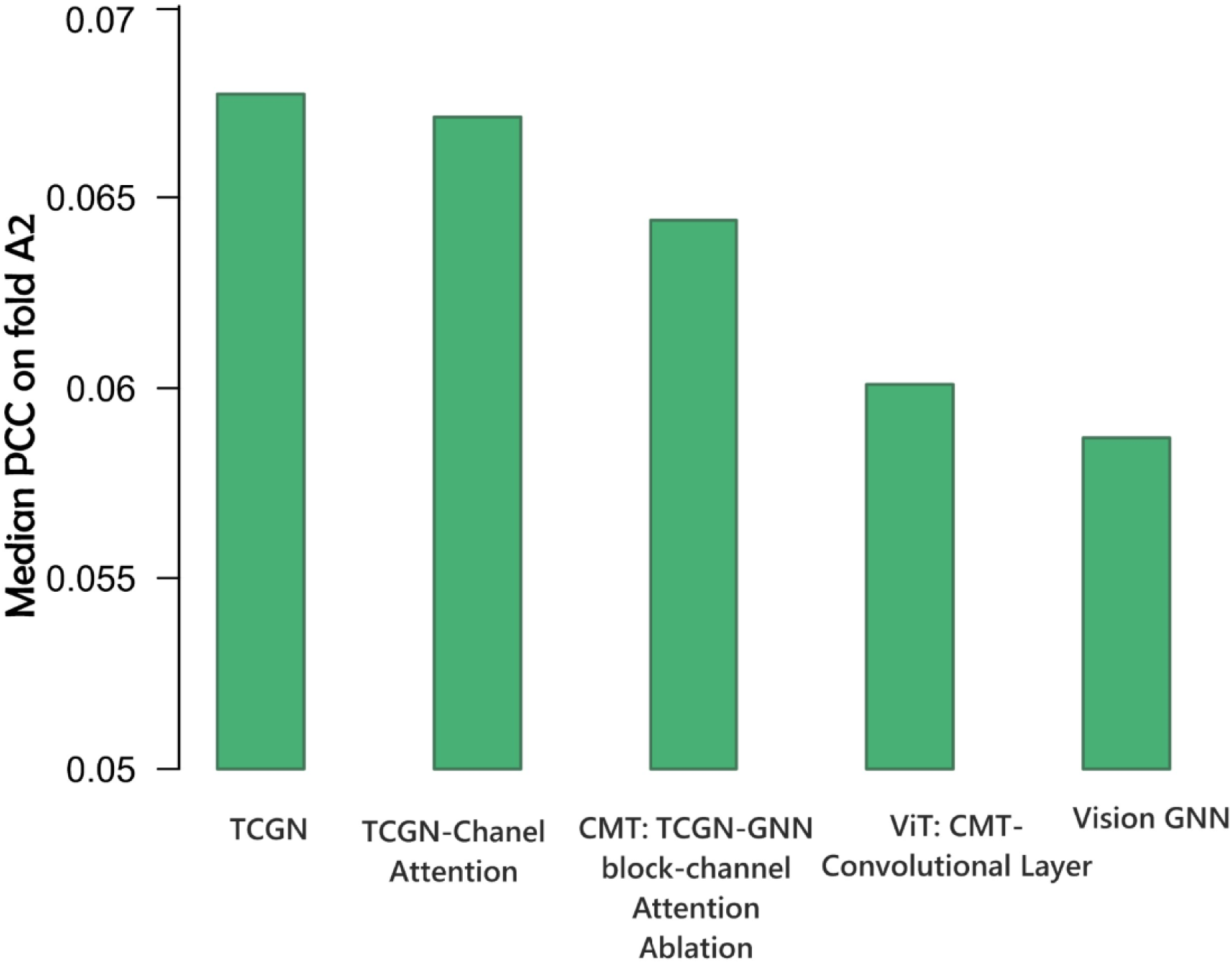
The ablation study on our model for examining the contribution of each block to the whole model.

### 3.3 Visualization of the genes with top prediction accuracy

To explore what kinds of genes can be predicted only by the local histopathological image without coordinate information, we ranked all the selected 785 highly variable genes according to their mean p-values among 32 cross-validations, where the p-value was assessed under the null hypothesis that there is no positive correlation between the predicted gene expression and the observed gene expression. The top 4 genes that have the highest mean p-values by TCGN are CLDN4, GNAS, MYL12B, and SCD, 3 genes overlapping with the top 4 genes found by HisToGene (GNAS, MYL12B, FASN, and CLDN4), and another 3 genes overlapping with that found by Hist2ST (FN1, GANS, SCD, and MYL12B). The top 4 genes with the highest prediction accuracy are GNAS, ACTG1, FASN, and DDX5, 2 genes overlapping with the top 4 genes with the highest mean p-values, indicating that some genes can be predicted well using local histopathological images without global information. After selecting the top genes, we then visualized these genes through the Python package scanpy[27]. Same as Hist2ST, we chose the tissue section with the highest PCC of the corresponding gene for visualization for each of the four genes (**Figure 5**). We can see that for all 4 genes in the corresponding tissue section, TCGA achieved the highest prediction accuracy. Although TCGN and other methods did not share the same tissue sections in which PCC was the highest for some genes, the predictions at the top 4 genes by TCGN were more accurate than that of other methods. For example, the highest PCC for MYL12B is 0.657 in TCGN, 0.621 in Hist2ST, and 0.557 in HisToGene.

**Figure 5.**
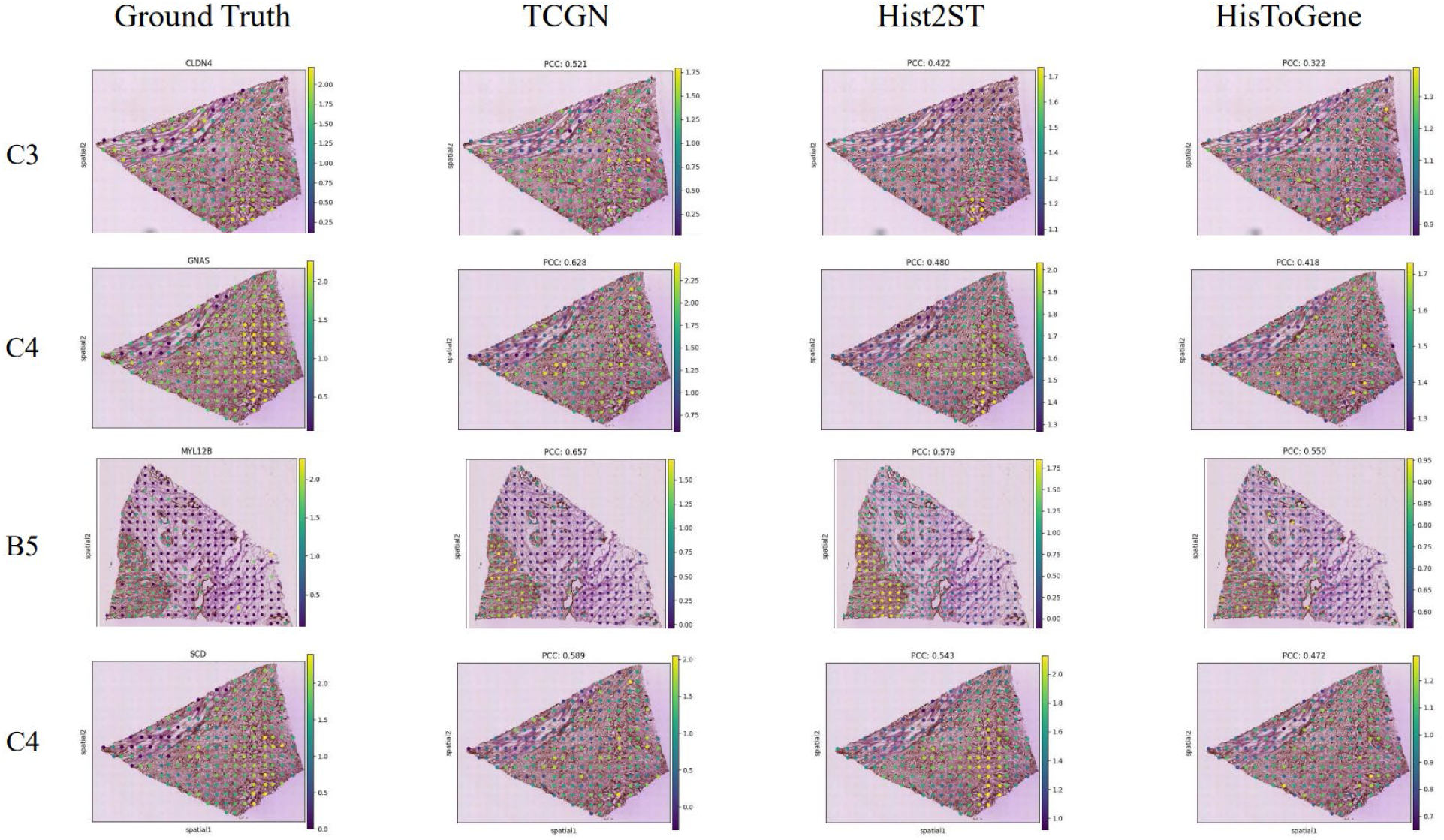
Visualization of the top 4 genes with the highest average p-value among the 32 tissue sections in the cross-validation. The p-value was generated by the hypothesis testing whose null hypothesis is there is no positive correlation between the predicted gene expression and the observed gene expression, and the tissue section with the highest PCC of the corresponding gene was selected for visualization for the 4 genes.

### 3.4 Functional gene analysis

To understand what kinds of genes can be predicted by local histopathological image and why our deep learning model can predict them with relatively high accuracy, we analyzed the functions of the top 4 genes with the best prediction accuracy, the genes with mean p-values less than 0.05, and the genes with mean p-values less than 0.1.

The top 4 genes with the best prediction accuracy are related to breast cancer and are potential biomarkers for cancer. CLDN4 is a member of the tight junction proteins family and has been found to be abnormally expressed in breast cancer and other cancer types[41] and contribute to tumor invasion and tumor cell migration[42]. It is also a potential therapeutic target and indicator of malignant phenotype[43]. GNAS has been reported by several studies to promote cancer progression[44] and is a transcriptional biomarker[45]. Over-expression of the GNAS gene can activate the PI3K/AKT/Snail1/E-cadherin pathway, resulting in breast cancer cell proliferation and migration and reducing overall survival[46]. MYL12B is related to cell proliferation[47]. More interestingly, one study has demonstrated that phosphorylation of MYL12 can change cellular shapes in Cochlear Hair Cells[48], which is similar to our findings and may indicate that MYL12 can also be predicted in breast cancer cells. For stearoyl-CoA desaturase (SCD), it has been reported that this gene is highly correlated with ferroptosis and is overexpressed in cancer cells[49], indicating that our model can be a potential method for the examination of the new therapeutic strategy concerning ferroptosis[50].

Besides the top 4 genes, we also quantified what genes can be predicted with high accuracy in the tissue sections based on the 32 cross-validations and performed a gene ontology (GO) enrichment analysis on these genes. The enriched terms for the genes with mean p-values less than 0.05 and 0.1 were shown in **Figure 6**. Most of the enriched GO terms are related to NADH-related enzymes and the MHC proteins’ function, which are similar to the results in Hist2ST where the NADH dehydrogenase activity was the top enriched pathway and the results in ImageCCA where genes involved in immune response and cell-cell signaling (corresponds to the MHC protein function in our study) were correlated with the morphology[2].

**Figure 6.**
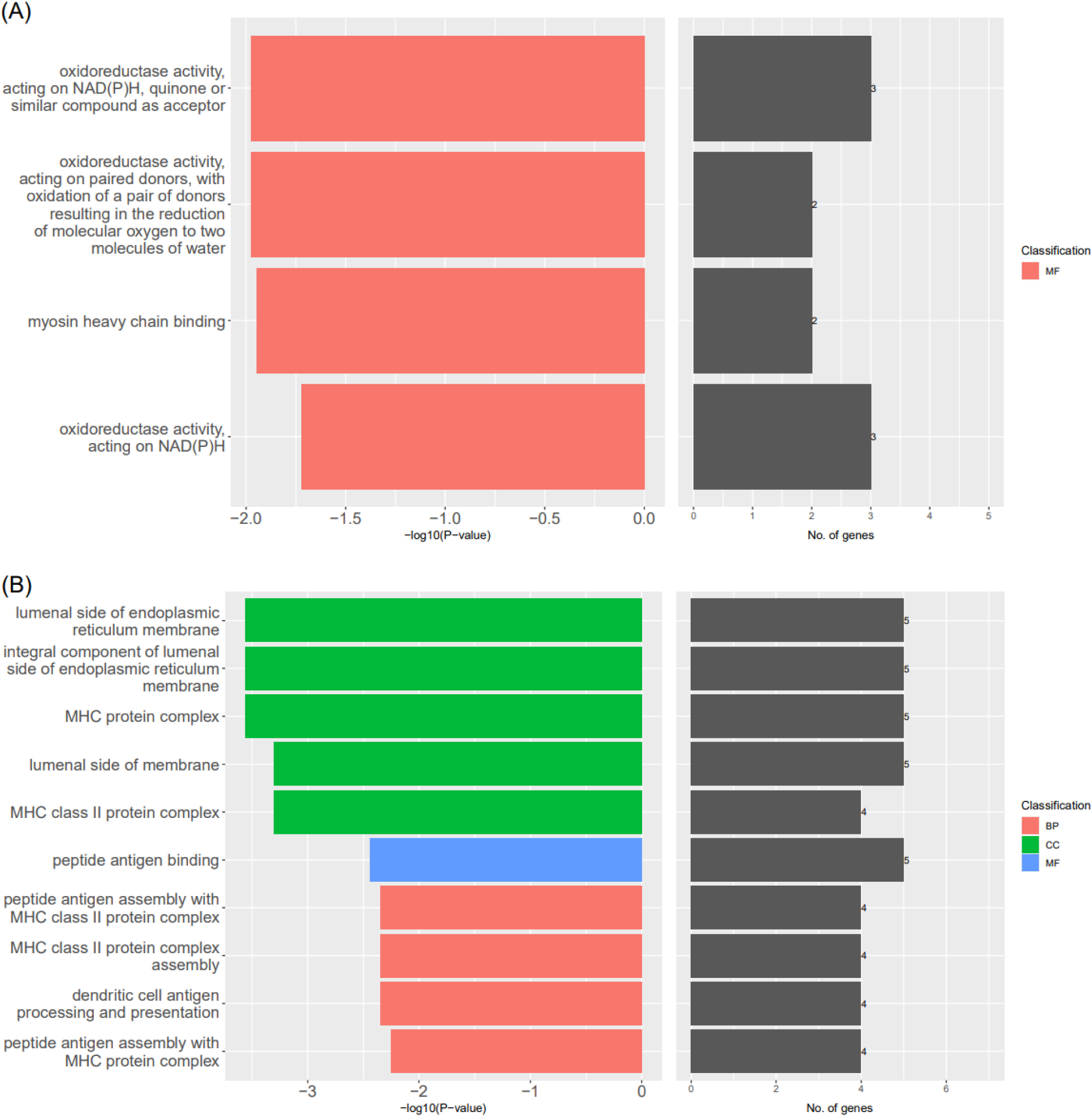
**A.** The enriched GO terms for the genes with the mean p-values less than 0.05 among the tissue sections in the 32 cross-validations, where BP, MF, and CC donate biological process, molecular function, and cellular components, respectively. **B.** The top 10 enriched GO terms for the genes with the mean p-values less than 0.1 among the tissue sections in the 32 cross-validations.

### 3.5 Interpretability and super-resolution gene expression

We additionally performed a Grad-CAM analysis to visualize the importance of every pixel in the input image in order to interpret our deep-learning model. Unlike previous studies that used all the spots in a tissue section and their coordinate information as the input, which can be potentially difficult to perform an interpretability analysis, TCGN is a general histopathological image analysis backbone, so we can analyze which parts in the image are important for gene expression predictions. Grad-CAM is a gradient-based localization method that creates coarse heatmaps from neurons whose gradients have a favorable impact on a class of interest in order to provide visual explanations for classification and regression tasks on images, and has been widely used in the interpretability of biomedical image analysis[51]. In this work, we randomly picked an image from the tissue section A2 and generated the Grad-CAM graph for the top 4 genes with the best prediction accuracy. **Figure 7** indicated how each pixel in the image contributed to the overall gene expression level in the spot and is also the pixel-level super-resolution gene expression in the spot.

**Figure 7.**
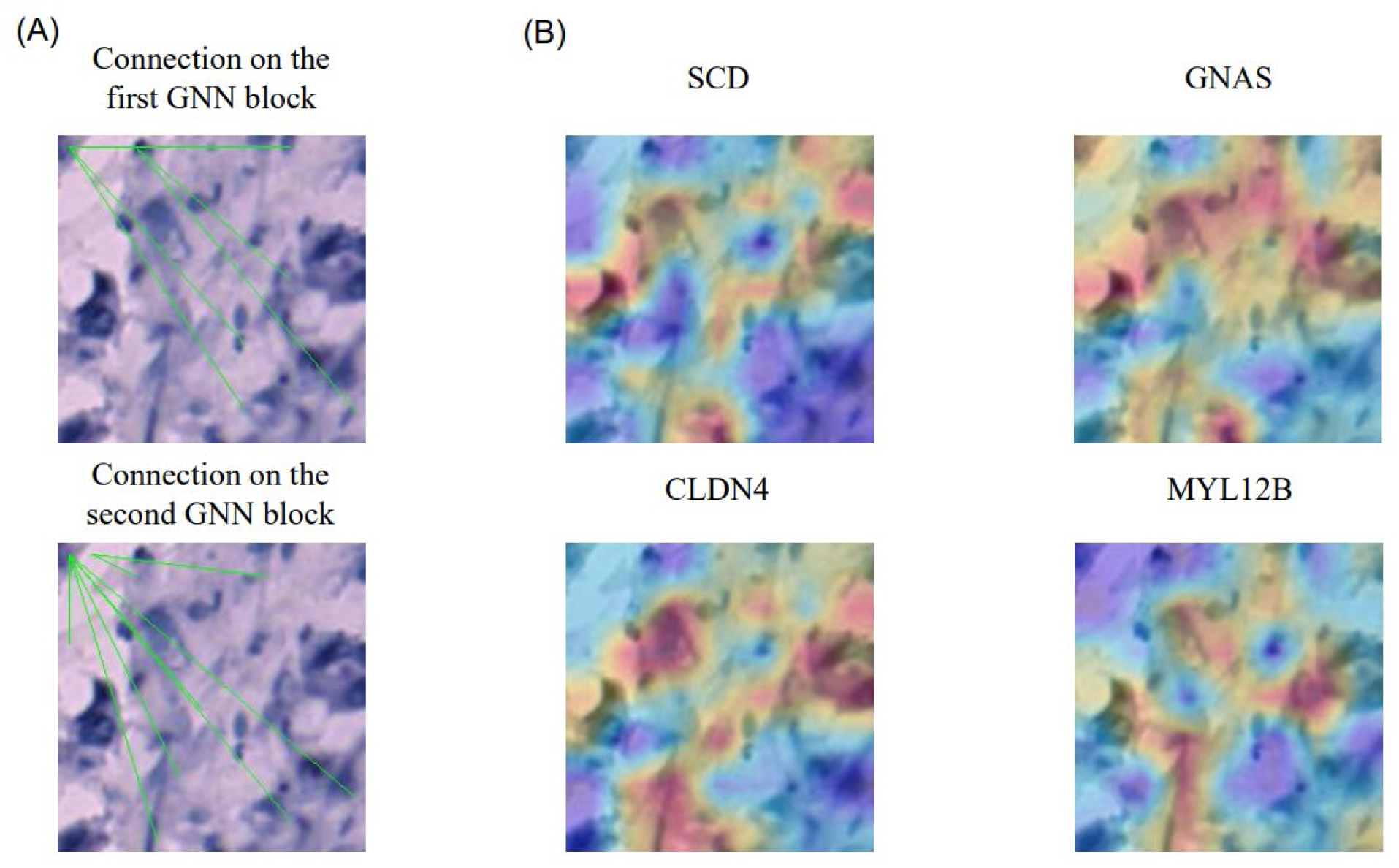
Demonstration of our model’s interpretability. **A.** The first 10 connections of the first and second left top patches generated by the connection analyzer. **B.** The Grad-CAM graph of one spot in tissue section A2 with the classes of interest which are the 4 genes with the top prediction accuracy.

In addition, we also analyzed what graph-structured features were extracted by the connection analyzer and learned by our model. Considering that there are number_of_patches × topk edges in one image, which was too numerous to demonstrate, we selected the first 10 connections of the first and second left top patches demonstrated in Figure 7. As shown in the graph, our connection analyzer can capture the relationships between cell nuclei and cytoplasm with certain morphology.

## 4. Discussion

### 4.1 How does the deep-learning model predict gene expression?

In this work, we demonstrated that genes can be accurately predicted even without global information if we can make full use of the features in the local spot image. From our perspective, we believe that gene expression can be predicted because their expression level is determined by cell type, the shape of cytoplasm around different cell nuclei, and cell-cell interactions (cell organizations)[2, 52], where all these features can be extracted by different components in our model.

#### 4.1.1 Identify different cell types to make gene expression prediction

There are numerous works that have proven the power of CNN in extracting the cell features from histopathological images and classifying cells, because CNN is very good at extracting local features of the input image. This can be briefly summarized by all kinds of UNet[53] that used the convolutional layers to extract cell features and utilize these features to segment and classify the cells[54, 55] in histopathological images. Also, many works concerning single-cell RNA sequencing have proved that different cell types have gene expression patterns that are largely different from each other[56]. Since the CNN can identify the cell types, it is possible that the CNN block in our model can analyze the potential cell types in the input and use them to infer the gene expression levels in these cell types. Then the overall gene expression level can be predicted through the average pooling layer of our model which can aggregate the result from different parts of the processed image (feature maps). To take the Grad-CAM graph of the gene GNAS in Figure 7 as an example, we can learn from the figure that the 3 cells with abnormal cell nucleus sizes that are close to each other contribute most to the overall expression level of GNAS in this spot. And it is possible that these 3 cells belong to the cancer cell that has a larger cell nucleus than normal cells. The distance and the number of these possible cancer cells may indicate the occurrence of tumor migration and tumor cell proliferation thus indicating a high level of GNAS expression.

#### 4.1.2 Detect abnormal cytoplasm to infer abnormal gene expressions

There have been many works reported that abnormal gene expression can be reflected in organelles and cell membranes within the cytoplasm[2, 48]. It is also reported by ImageCCA[2] that the genes that can be predicted by the histopathological images are related to cell membrane functions such as cell adhesion and the cell component - intracellular organelle. And this is reasonable not only because the shape of the cytoplasm can help identify the cell types, but also because the texture of the cytoplasm may be an indicator of gene expression. For example, the GNAS gene is related to the cell adhesion function, and the cancer cells are less adhere to each other compared to normal cells, so a set of cells with large cell nuclei and abnormal cytoplasm texture may indicate the cell type of cancer cell and these cells’ abnormal expression of GNAS. And this is the case in Figure 7 where the regions that contain cells with large cell nuclei and blurry cytoplasm contribute most to the GNAS gene’s expression prediction. In addition, the size of the cytoplasm can also reflect the activity and the growth condition of the cell and the image region. This may be the reason why the genes with top prediction accuracy are enriched in NAD(P)H functions and oxidoreductase activity (Figure 6), which is a crucial part of respiration and may be able to measure the cell activity in this image region.

In our model, with the help of CNN and transformer encoder, we can extract different cytoplasm texture features and pay attention to the features among them that contribute most to the final gene expression prediction task. Also, the pyramid structure of our deep learning model makes it possible for the model to learn more high-level and global features such as shape features and cell organization features in deeper layers, which let our model be able to make full use of the cytoplasm features to predict the genes accurately.

#### 4.1.3 Use cell-cell interactions to adjust morphological features and make predictions

Unlike previous GNN-related histopathology studies[23, 33, 57], the GNN block in our model can analyze the interaction such as connections between cell nuclei and cytoplasm (Figure 7), other than just the connection between cell nuclei and cell nuclei. And the complex cell nuclei segmentation operation is not needed.

The cell-cell interaction features contribute much to the overall gene expression levels in the spot[52] and can be reflected in the cell organization that contains the cell nucleus and cytoplasm. The cell-cell interactions not only directly influence the gene expression level through different weights of graph features in the model, but also contribute to the gene expression prediction task by adjusting the contribution of every cell or every patch in the input local spot image. For the direct effect of the graph features, a successful application is that the Pathomic Fusion[23] used the graph neural network to extract the features of spatial cell graphs to make the prognostic predictions with a high C-index. And a biological example is that there will be MHC protein-related signal transduction when there are cancer cells and immune cells and there is a connection graph between these cells in the input image. And this may be the reason why the genes with top prediction accuracy are enriched in MHC proteins’ functions. For the “adjusting” effect, it is easy to understand that even for the same type of cell, the expression of this cell would also change according to the environment this cell is in. So in our model, we used the channel attention block to simulate this “adjusting”, where the interactions between features generated by the GNN block and other blocks are analyzed by the attention mechanism.

Our model can learn cell organization features efficiently with the help of the GNN block, the transformer encoder, and the pyramid structure. For the GNN block in our model, there have been researchers reported that the cell-cell interactions can be extracted by the graph neural network[23]. In addition, since the transformer encoder is a special form of GNN[35], the CMT block in our model can also extract the graph-structure features that represent the cell-cell interactions. And as illustrated above, the pyramid structure of our model also enables our model to extract the cell organization features at different complex levels. In addition, compared to previous works that used the segmented cell nucleus as nodes and the distance between the nucleus as edges, our model can learn the graph-structure features better because we use the neural network to learn different edge features and we treat both cell nucleus and cytoplasm as nodes instead of only the cell nucleus. Another advantage of our model is that we also do not need complex cell segmentation to build the spatial cell graph thus making the analysis process more simple and more accurate.

### 4.2 What genes can be reflected on histopathological images and why they can be predicted?

As shown in Figure 7, we can learn that the genes with high prediction accuracy mainly correlate with MHC protein functions, reticulum membrane, and oxidoreductase activity. So we speculate that the genes that can be predicted by local histopathological images should have an indirect or direct impact on the shape and texture of the cell nucleus and cytoplasm, expressed differently in different regions or cell types, and can influence the organization and the survival possibility of cells.

The indirect influence refers to the condition where the deep learning model can first detect the abnormal morphology, then the genes that have an indirect impact on and a close relationship with cells’ features can be detected by the deep learning model. And this is the case for the genes related to the MHC proteins’ functions and reticulum membrane. The MHC proteins, which are closely related to immunological recognition and antigen assembly, may indirectly influence some special cells’ survival rate and change the morphology of the region. It is reported that the abnormal expression of the MHC protein influences immune cells’ recognition of tumor cells, thus increasing the spread and proliferation of tumor cells[58]. And since tumor cells have special morphology and can be detected by our deep learning model, then the regions where there are many cancer cells will be recognized as having abnormal MHC proteins-related gene expressions. This is also the case for genes related to the reticulum membrane, where there have been researchers reported that dysregulation of endoplasmic reticulum morphology is tightly linked to neurologic disorders and cancer[59].

For the direct influence on morphology, the genes related to the reticulum membrane, and oxidoreductase activity may directly change cells’ morphology by affecting the growth, development, and functioning process of cells. For the reticulum membrane, the disorder of the reticulum membrane may affect the function of the cell membrane as the reticulum is responsible for the processing of membrane proteins. And the malfunction of membrane proteins will affect the cell-cell adhesion, resulting in shape changes in the cytoplasm. Moreover, the abnormal reticulum can also be reflected in the features of the cytoplasm such as the degree of image blur. As for the oxidoreductase activity, it is known that cancer cells normally have a much more intensive metabolism and a high oxidoreductase activity, and the abnormal oxidoreductase activity will make the cancer cell appear differently in terms such as its nucleus shape. Then the shape feature can be recognized by our deep learning model and let it make an accurate expression prediction.

### 4.3 The potential of our TCGN architecture

As a general and powerful computer vision backbone with simple input for histopathological image analysis, our model can predict gene expressions and biomarkers accurately from only the local histopathological images, and this makes it a potentially powerful method for other clinical tasks concerning histopathological images such as prognostic prediction and precision medicine guidance.

For prognostic prediction, where many works have shown that applying deep learning to histopathology-based prognosis analysis can largely increase its speed and accuracy[60], several works in this field simply used the convolutional neural networks to encode the local histopathological image and aggregate or make statistical analysis on them[61]. But since we proved that using the integration of GNN, transformer encoder, and CNN can make a better analysis of histopathological images compared to only the CNN, applying our model to these tasks may make the results better. And this can also be illustrated by the genes with the top prediction accuracy in our study, where most of them are closely related to breast cancer and lead to a better or worse prognosis.

As for precision medicine guidance, the accurate gene expression predictions in our model have proved the TCGN’s power in detecting biomarkers and classifying disease subtypes. For example, the high prediction accuracy of the gene SCD indicates that our model may be able to help test whether further ferroptosis-related chemo-immunotherapy[62] is needed. Another example is that the RHOB gene, which is the gene with the top 6 prediction accuracy, is an important indicator for classifying different breast cancer subtypes[63]. So it is possible that we can simply analyze patients’ histopathological images to know patients’ exact cancer subtypes and provide precision medicine guidance to them.

In addition, since our model is interpretable and we can make the Grad-CAM graph, our model can also analyze the pixel-level super-resolution gene expression using the histopathological images.

### 4.4 Multi-task learning: median PCC V.S. MSE

The loss function in our model and the previous 3 works were all the MSE loss, but it seems that there is still much to improve for the loss function. The reason for this statement is that there is no positive correlation between the median PCC on the validation set and the MSE loss on the validation set. That means the model chooses to increase the prediction accuracy of some genes while decreasing the prediction accuracy of other genes in order to make the sum of all genes’ prediction errors the minimum. So in our work, for the purpose that the model can predict more genes accurately, we choose to save the model’s parameters when the median PCC on the validation set reaches its peak.

As shown in **Figure 8**, in the first 4 epochs, the MSE loss keeps decreasing while the median PCC keeps increasing. And then in epoch 5 to epoch 15, the MSE does not change much, and the median PCC fluctuates between 0.06 and 0.07. These phenomena are predictable so far. But in epochs larger than 16, the MSE loss continues to decrease, but the median PCC also decreases.

**Figure 8.**
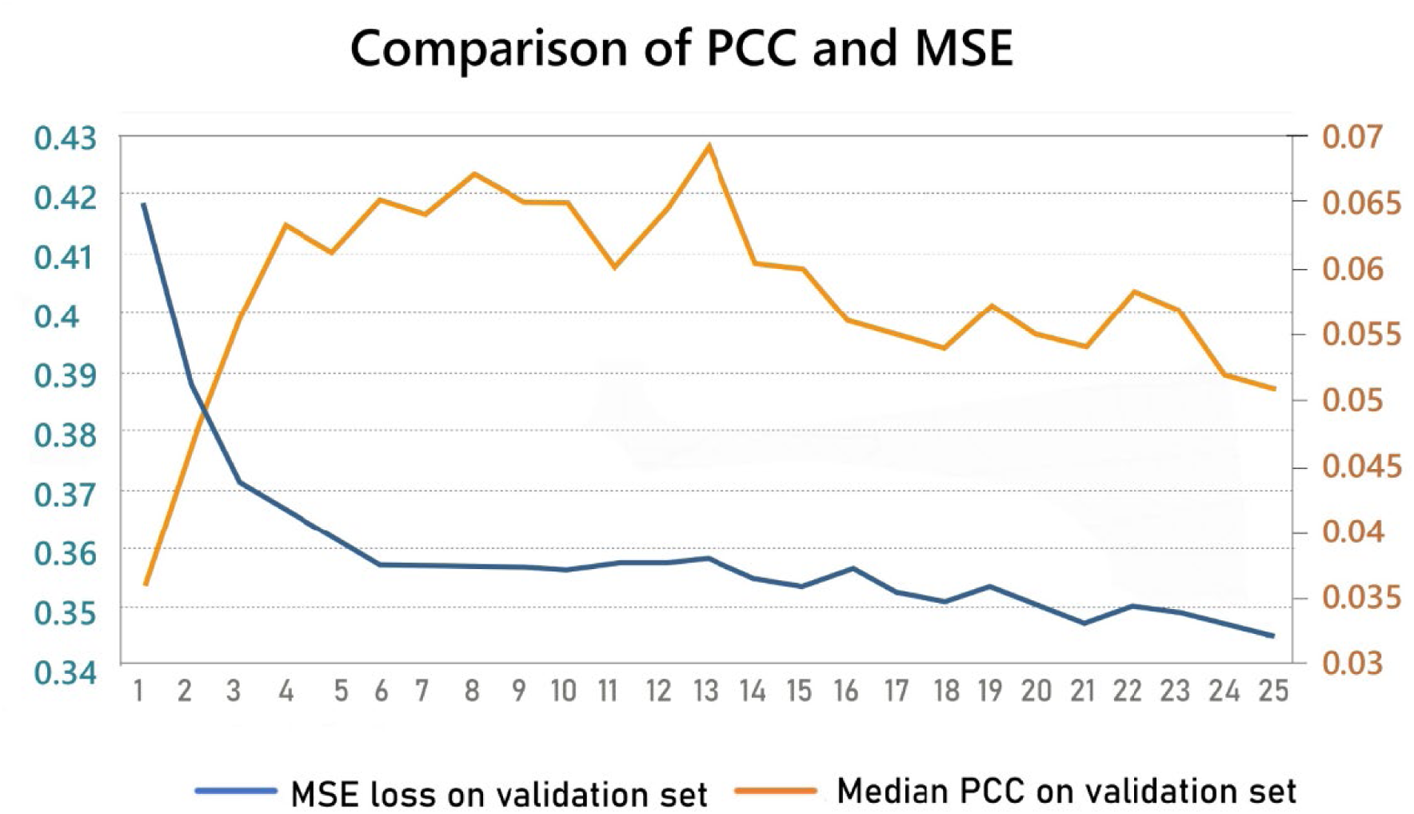
The changes of the MSE loss and the median PCC on the validation tissue section A2 as the epoch increases.

The reason for the non-positive correlation is that every gene contributes the same to the final loss. And since all the predicted genes shared the same backbone/features extractor and the model used the gradient descend method to be optimized, it is very possible that the features of the histopathological image that is important for most genes will be extracted and the features that are only helpful to fewer genes’ prediction will be ignored. In our view, giving different genes different weights for the final loss and changing the weight according to the prediction performance of each task in each epoch may predict every gene more accurately, but this requires better multi-task deep learning theory in the future.

### 4.5 Use adam optimizer instead of the stochastic gradient descent

Although the prediction accuracy of ST-Net was very low when using the hyperparameters and the optimizer (stochastic gradient descent) provided in the corresponding paper[3], if we change the optimizer to the adam optimizer, the mean of all genes’ mean PCC of ST-Net would be largely increased to 94% of that of HistToGene. And this is about 10 times higher than the accuracy when using the stochastic gradient descent for ST-Net. This phenomenon may be caused by the situation where different patches indicate different relationships between the genotypes and the phenotypes, and these relationships have something in common but have some kinds of noise in each of them. And this is common sense in biology, where there is a lot of randomness and noise in the biological experiments, but people can always conclude something from these experiments. So when using the stochastic gradient descent, the model cannot learn a stable relationship between the genotypes and the phenotypes because of the negative influence brought by the noise. But when we use the adam optimizer to utilize the gradient of previous training steps, the gradient descent would be more stable and less vulnerable to the noise, thus having a better performance. This may also be the reason why the Hist2ST and HisToGene that use all the spots as input outperformed the ST-Net. Because treating all the patches as input makes the model learn the sum of all the relationships reflected on every patch, the negative influence of the noise would then be smaller.

### 4.6 Limitations

Despite the outstanding performance of our model in predicting gene expression from only local images, our model still suffers from the following 3 drawbacks, and due to objective conditions, our work can be improved by using more breast cancer spatial transcriptomics data to further validate our model.

The first shortcoming of our model is in the top_k selection step in the building graph block, where we simply add the number_of_patches × topk by 1 if there is more than one value ranking the top *number*_*ooo*_*patches* × *topk* to ensure the number of selected edges to be the same within one batch. But this would cause the number of edges to differ from that of another batch, and let the model deal with unequal and confusing input thus lowering the final prediction accuracy. Secondly, some augmentation methods such as self-distillation can be applied to our model to eliminate the error caused by the random initialization of our model and further improve the prediction accuracy. And thirdly, it is possible that pretraining our model on the ImageNet[64] can improve the prediction accuracy since our model only requires a 224 × 224 input like other computer vision backbones. And this can also solve the problem of the relatively small amount of biomedical image data.

## 5. Conclusion

In this paper, we proposed TCGN, a powerful deep-learning model to predict genes from only the local histopathological image of breast cancer patients. Our model consists of convolutional layers, the transformer encoder, and the graph neural networks, and to the best of our knowledge, is the first work to integrate these blocks in a general computer vision backbone for histopathological image analysis. Compared to previous studies, our work achieved a higher prediction accuracy with a smaller model size without the global information. And compared to the counterpart of our model with the same input type, our model achieved about 10 times higher gene prediction accuracy. We also analyzed the interpretability of our model, and we found that our deep-learning model can extract the features concerning special cell types, cytoplasm features, and cell organizations to predict gene expression from histopathological images. And for the genes with high prediction accuracy, we found that the genes with high prediction accuracy mainly correlate with MHC protein functions, reticulum membrane, and oxidoreductase activity, which have an indirect or direct impact on the shape and texture of the cell nucleus and cytoplasm, express differently in different regions or cell types and can influence the organization and the survival possibility of cells. Besides predicting gene expressions, our model may also be applied to many other domains like prognostic prediction and precision medicine guidance.

## 6. Code available

The code for the TCGN model is available on GitHub.

## 7. Data available

The human HER2-positive breast tumor spatial transcriptomics dataset we used in this paper is available at https://github.com/almaan/her2st/.

## 9. Author Contributions

X. X. participated in the data acquisition, performed the model design and the statistical analysis, and drafted the manuscript. Y.K. participated in its design and drafted the manuscript. Z.W. supervised the biostatistics analysis, and all aspects of the study were supervised by H.L. and Y. K. All authors read and approved the final manuscript.

## 11 Acknowledgment

The computations in this paper were run on the π 2.0 cluster supported by the Center for High-Performance Computing at Shanghai Jiao Tong University.

## 13. Vitae

Xiao xiao is an undergraduate student at Shanghai Jiao Tong University and will be a PH.D. student at Yale this year in 2023 Fall. He is interested in biological information research and deep learning modeling.

Yan Kong is a postdoc researcher at SJTU-Yale Joint Center for Biostatistics and Data Science, Shanghai Jiao Tong University. Her research focuses on new strategies for multi-modal data fusion and the applications of deep learning methods on medical images.

Zuoheng Wang is a professor in the Department of Biostatistics at Yale University. Her research focuses on biomedical longitudinal data modeling and biostatistical theory studies.

Hui Lu is a professor at Shanghai Jiao Tong University. His research focuses on statistical modeling and building large pre-trained deep-learning models.

